# ABCB-mediated auxin transport in outer root tissues regulates lateral root spacing in *Arabidopsis*

**DOI:** 10.1101/2020.07.22.206300

**Authors:** Jian Chen, Yangjie Hu, Pengchao Hao, Yuqin Zhang, Ohad Roth, Maria F. Njo, Lieven Sterck, Yun Hu, Yunde Zhao, Markus Geisler, Eilon Shani, Tom Beeckman, Steffen Vanneste

**Affiliations:** Department of Plant Biotechnology and Bioinformatics, Ghent University, 9052 Ghent, Belgium; Center for Plant Systems Biology, VIB, 9052 Ghent, Belgium; School of Plant Sciences and Food Security, Tel-Aviv University, 69978 Tel-Aviv, Israel; Department of Biology, University of Fribourg, CH-1700 Fribourg, Switzerland; Section of Cell and Developmental Biology, University of California San Diego, La Jolla, CA, USA; Lab of Plant Growth Analysis, Ghent University Global Campus, Incheon 21985, Republic of Korea

## Abstract

Root branching is an important strategy to explore efficiently large volumes of soil. To economize this process, lateral roots (LR) are formed along the growing root at discrete positions that are instructed by oscillating auxin signals derived from the lateral root cap (LRC). This assumes that auxin moves from the LRC across multiple layers to accumulate in the pericycle. Here, we identified, using gene silencing and CRISPR based approaches, a group of five genetically linked, closely related ABCBs that control LR spacing by modulating the amplitude of the auxin oscillation. The transporters localize to the plasma membrane and reveal significant auxin export activity. These ABCBs are mainly expressed in the LRC and epidermis where they contribute to auxin transport towards the root oscillation zone. Our findings highlight the importance of auxin transport in the outer tissues of the root meristem to regulate LR spacing.

## Introduction

The root system of plants is of vital importance for their growth and survival as it anchors the plant in the soil and is required for the uptake of water and nutrients and symbiotic interactions. The complexity of root systems can be easily expanded by LR branching according to environmentally imposed limitations and stimuli^1^. LR development is a multistep process occurring over a long time, involving coordinated signaling across several tissues^2^. The plant hormone auxin is a key regulator of many organogenetic events in plants^3^. Its local accumulation triggers dramatic, preprogrammed transcriptional changes that are associated with the progression of the developmental program^3^. This is also the case for LR development, where auxin accumulation defines the spacing of prebranch sites along the primary root, and thus root architecture complexity^4, 5, 6, 7^. Therefore, plants have established intricate mechanisms to control auxin distribution within tissues^8, 9,^ which can be adjusted according to the developmental stage, hormonal and environmental signals^1^. Under normal conditions, the initial decision for LR initiation is not made where low-sensitive auxin signaling output reporters in pericycle cells are observed, but rather in a zone more proximal to the meristem^6, 10^. In this zone, called the oscillation zone (OZ), oscillatory gene expression was reported to correlate with the activity of the sensitive auxin signaling output reporter *DR5::LUC*^5^. This periodic auxin signaling selects a subset of cells, together denominated as a prebranch site, to gain a higher competence to form a LR reflected in a maintained expression of the auxin output reporter *DR5::LUC*. ^5^. Indole-butyric acid (IBA) to indole-5-acetic acid (IAA) conversion in the LRC contributes to the amplitude of this oscillation^11, 12,^ and cyclic programmed cell death of the LRC contributes to the frequency of this oscillation^7^.

The prevailing model of auxin transport in the meristem can best be summarized as a reverse fountain of auxin flowing rootward through the vascular tissue and being redirected shootward through the outer layers of the meristem^13^. This outer shootward auxin flow is thought to rejoin the central rootward auxin flow (Fig. 1a). The radial inward movement of auxin released from LRC cells undergoing programmed cell death could then contribute to periodic peaks of *DR5:LUC* that are instructive for LR positioning^7^. The IAA^−^/H^+^ symporter AUX1^14^ represents the major component in the auxin uptake mechanism of the inward radial auxin transport route that controls prebranch site formation^6, 7^. However, the corresponding efflux components remain elusive. Currently, the auxin efflux component of the shootward auxin transport in the outer cell layers of the root is believed to be largely explained by the PIN2 auxin transporter^15,^ in conjunction with the ABCB-type auxin transporters, ABCB1 and ABCB19^16^. However, neither the *pin2* mutant, nor the *pin2,abcb1,abcb19* triple mutant showed reduced LR densities or reduced IBA-induced LR formation^7^. This suggests that our current model contains important inaccuracies or shortcomings at least at the level of the molecular identity of the auxin efflux machinery in the root meristem.

**Fig. 1:**
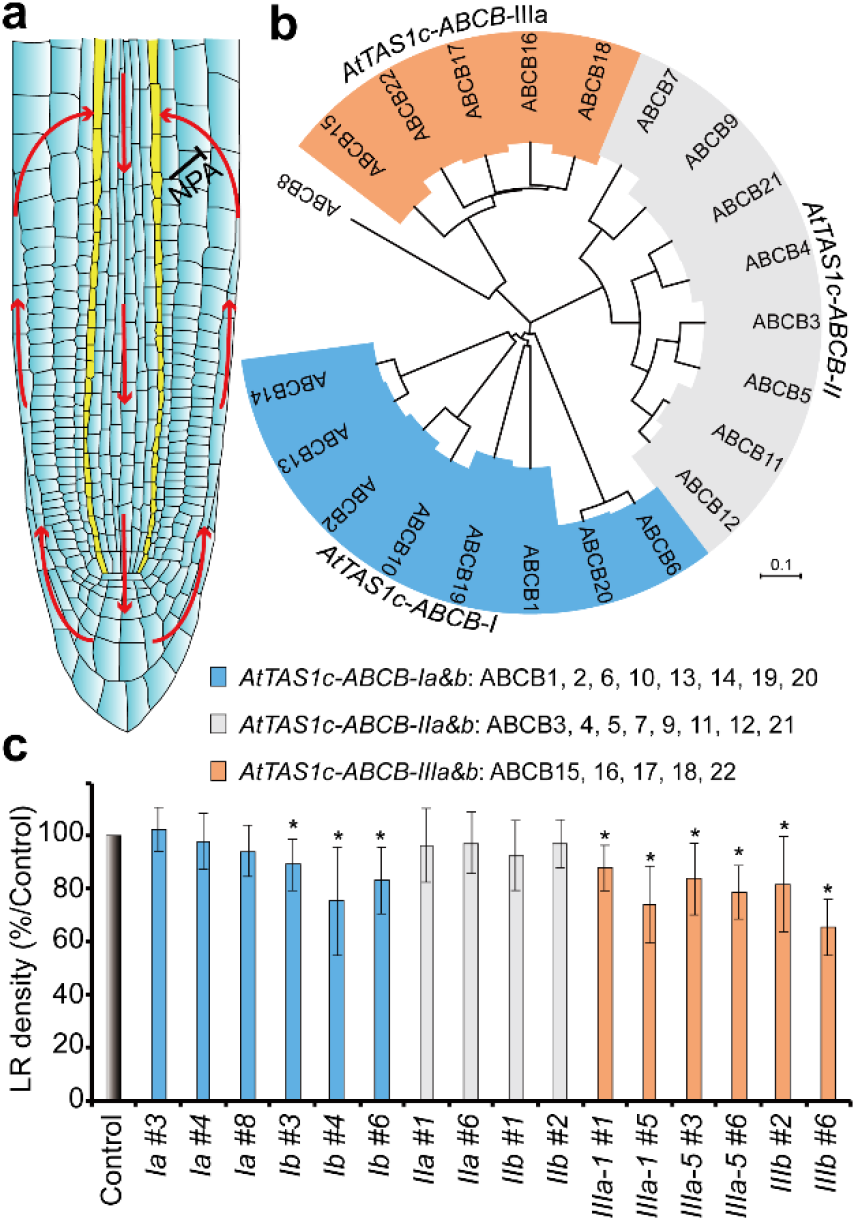
Phenotypic screen of the *ABCB* gene family for regulators of LR formation using *PIN2* driven syn-tasi-RNA-based gene-silencing. **a** Reverse fountain model of auxin transport in the *Arabidopsis* root apical meristem. Red arrows show the major auxin transport directions. The pericycle is indicated in yellow. **b** Phylogenetic tree of *ABCB* gene family. Subgroup genes targeted by different synthetic AtTAS1c constructs are colored blue (*AtTAS1c-ABCB-I*), grey (*AtTAS1c-ABCB-II*) and orange (*AtTAS1c-ABCB-III*). Syn-tasi-RNA constructs are driven by the *PIN2* promoter. **c** Quantification of the emerged lateral root density of 10-day-old *AtTAS1c* lines relative to WTs grown on the same plate in the initial phenotypic screen, shown are averages (±SD) of > 15 plants. WT controls are set to 100%, * indicates P < 0.05 by two-tailed Student’s t-test relative to WT. Colors correspond to different subgroups as indicated in **b**.

Here, we aimed at identifying auxin transporters in the outer tissues of the root that have a role in LR spacing. Therefore, we screened the *ABCB* transporter family via a tissue-specific gene-silencing approach. We found five closely related ABCBs that are required for LR spacing via modulation of the *DR5:LUC* oscillation amplitude. These ABCBs localize to the plasma membrane and transport auxin out of the cell. Their predominant expression in LRC and epidermis, and the LR defects in the knock-down/out lines is consistent with these ABCBs acting as effectors of auxin transport in the outer layers of the root meristem that instructs LR spacing.

## Results

### An uncharacterized cluster of five ABCBs controls LR density

Lateral root initiation and spacing depend on an intricate auxin transport mechanism in the root meristem that can be inhibited via auxin-transport inhibitors such as NPA and BUM^7, 17, 18^ (Fig. 1a). Because both inhibitors are thought to target, among others, the auxin-transporting ABCBs^18, 19, 20,^ we reasoned that members of the full-sized 22 ABCB protein family^21^ (Fig. 1b) are involved in auxin transport for LR spacing.

To screen the *ABCB* family for members that are potentially involved in inward radial auxin transport (between the LRC and the pericycle), we used synthetic trans-acting small-interfering RNAs (syn-tasiRNAs) in the *AtTAS1c* backbone^22^. Based on three syn-tasiRNAs per ABCB subgroup we could theoretically target 21 out of 22 ABCBs: subgroup I: ABCB1,B2,B6,B10,B13,B14,B19,B20 (*AtTAS1c-ABCB-I*), subgroup II: ABCB2,B3,B5,B7,B9,B11,B12,B21 (*AtTAS1c-ABCB-II*), and subgroup III: ABCB15,B16,B17,B18,B22 (*AtTAS1c-ABCB-III*) (Fig. 1b). For each subgroup, we designed two independent and distinct syn-tasiRNAs to account for variation in silencing efficiency and specificity (indicated as ‘a’ and ‘b’) and expressed them in the LRC, epidermis and cortex via the *PIN2* promoter^23^. We determined the LR density of at least two independent, homozygous, single locus lines per construct, relative to WT grown on the same plate, hence analyzing at least 4 independent lines per ABCB subgroup (Fig. 1c). One of the constructs targeting subgroup I ABCBs (*AtTAS1c-ABCB-Ib*) resulted in a significant reduction of LR density, and both constructs reduced the primary root length (Fig. 1c, Supplementary Fig. 1). None of the *AtTAS1c-ABCB-II* lines showed a significant change in LR density or root length. Interestingly, all tested (six) *AtTAS1c-ABCB-III* lines showed significant reductions in LR density and root length (Fig. 1c, Supplementary Fig. 1). We could validate the reduction in LR density of 4 independent *AtTAS1c-ABCB-III* lines, but not the reduction in root length, which was seen in the initial phenotypical screen (Supplementary Fig. 2). This highlights *ABCB15*, *B16*, *B17, B18* and *B22* as potential regulators of LR development. In order to test whether single knockouts would be sufficient to express a LR phenotype, we obtained T-DNA insertion lines in these *ABCBs*. However, none of them showed a defect in root growth and LR formation (Supplementary Fig. 3), indicating a strong functional redundancy as described for other ABCB family members^24, 25, 26, 27, 28^.

Importantly, all members of the subgroup are clustered within 57.6 kb on chromosome 3, limiting the generation of multi-knockouts by crossing T-DNA lines. Therefore, to validate the observed *AtTAS1c-ABCB-III* phenotypes, we followed two independent knock-down/out approaches. On the one hand, we identified, within a collection of artificial microRNA (amiRNA) lines^26,^ *pro35S::amiR-2572* (named *amiR-2572*),that targets several of the same ABCBs as the syn-tasi constructs (*ABCB16*, *ABCB17*, *ABCB22*) (Supplementary Fig. 4a). On the other hand, we generated an *abcb*^*b15,b16,b17,b18,b22*^ mutant via genome editing using 4 sgRNAs, each targeting multiple members of this *ABCB* subgroup (Supplementary Fig. 4b). We identified one line, named *b15-b22*^*CRISPR*,^ in which we could not amplify a 3502bp fragment of *ABCB15*, a 1121 bp fragment of *ABCB16*, a 1520 bp fragment of *ABCB17* and which had a deletion in *ABCB18* and a single bp insertion in *ABCB22* that causes a premature stop codon, thus likely being a null mutant in these genes (Supplementary Fig. 4c,d). Using Nanostring-based mRNA quantification, we could, however, not reliably detect transcriptional downregulation of any of the targeted *ABCB*s in our knock-out/down lines, with the exception of *ABCB15*, being significantly downregulated in *b15-b22*^*CRISPR*^ (Supplementary Fig. 4a). Additionally, the non-targeted *ABCB4* and *ABCB19* were respectively up- and downregulated in *b15-b22*^*CRISPR*^ (Supplementary Fig. 4a) indicating complex interactions among auxin-transporting ABCBs, calling for caution in interpreting observed phenotypes.

Both *amiR-2572* and *b15-b22*^*CRISPR*^ showed a strong reduction in primary root length (Fig. 2b,c,) and LR density (Fig. 2b,d). Moreover, both lines had a smaller rosette size in soil and *b15-b22*^*CRISPR*^ had reduced fertility (Supplementary Fig. 4e,f). This suggests that the non-tissue-specific interference with the expression of these ABCBs results in pleiotropic phenotypes. In another genome-editing approach we generated a 54kbp deletion mutant in which the entire region was deleted, whilst retaining a chimeric fragment of both ABCB15 and ABCB22 (Supplementary Fig 5a,b). In contrast to *b15-b22*^*CRISPR*,^ this large fragment deletion (*ldf-A7*) mutant did not display any obvious phenotypes (Supplementary Fig 5c-f,), suggesting the induction of a compensation mechanism, eg. similar to the recently described interallelic complementation^29,^ or via a compensatory (non)-transcriptional activation of other transport mechanisms. Despite the unresolved issue with the 54 kb deletion mutant, we argue that 3 independent knock-down constructs and one CRISPR knock-out showing similar LR phenotypes represent compelling evidence for the involvement of the subgroup III ABCBs in LR spacing.

**Fig. 2:**
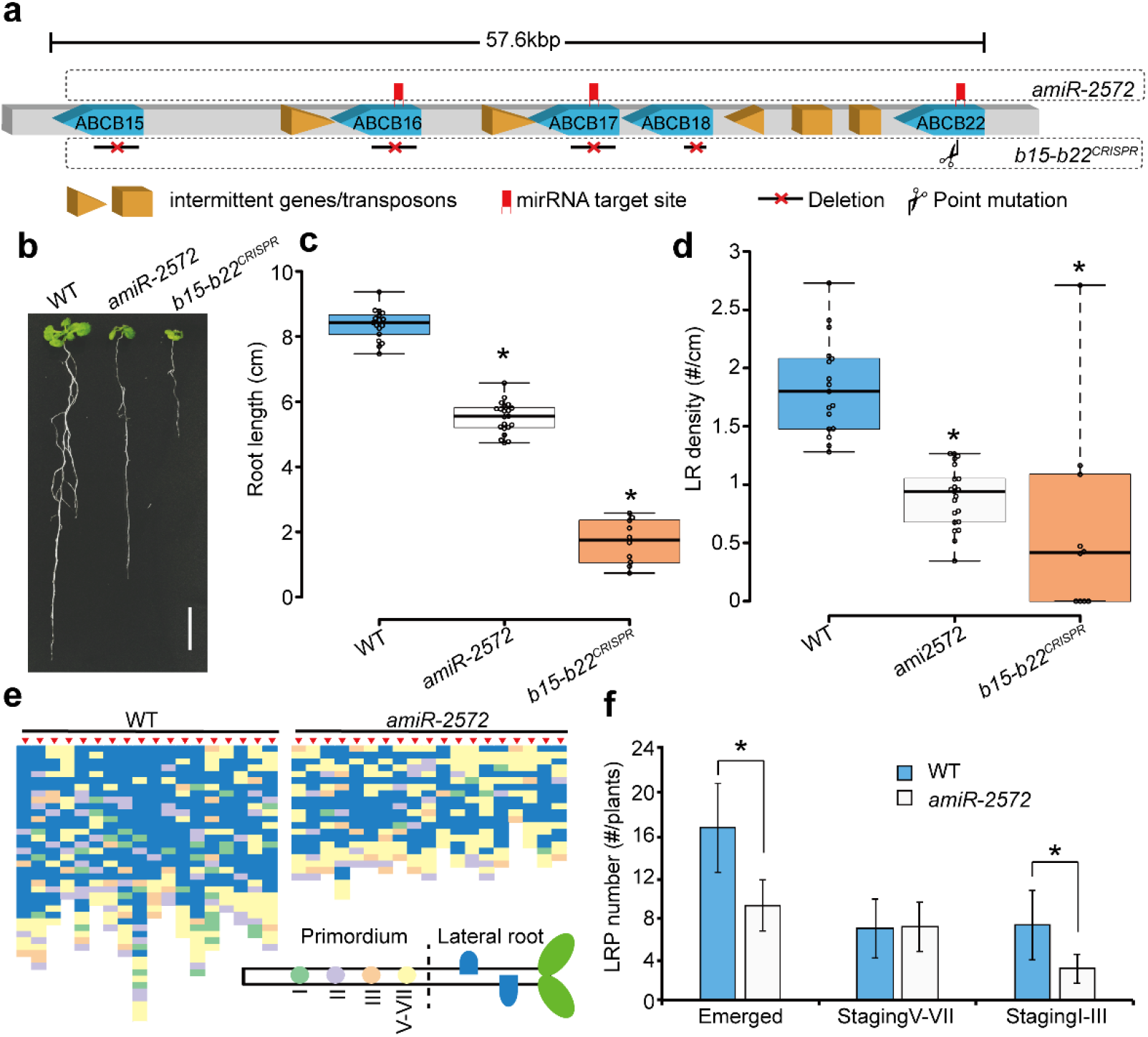
amiRNA and CRISPR confirm the importance of subgroup III ABCBs in LR development. **a** Organization of subgroup III ABCBs on the chromosome. All *ABCB* coding-regions are highlighted in blue, intermittent genes and transposons are highlighted in orange. *amiR-2572* target sites in the respective *ABCB*s are indicated with red marker in the square upper. The deletion within *ABCB15, 16, 17, 18* and the point mutation site of *ABCB22* in *b15-b22^CRISPR^* are indicated in the square below. **b** Macroscopic seedling phenotype of 15-day-old *amiR-2572* and *b15-b22^CRISPR^* compared to WT. Scale bar is 1 cm. **c, d** Boxplots showing the quantification of emerged lateral root number (**c**) and root length (**d**) in 15-day-old WT, *amiR-2572* lines and *b15-b22^CRISPR^* mutant seedlings. Center lines show the medians; box limits indicate the 25th and 75th percentiles as determined by R software; whiskers extend 1.5 times the interquartile range from the 25th and 75th percentiles, outliers are represented by dots; data points are plotted as open circles. n = 17 (WT), 21 (*amiR-2572*), 10 (*b15-b22^CRISPR^*), * indicates P < 0.05 (two-tailed Student’s t-test). **e** Schematic representation of the distribution of emerged LRs and LRP along the primary root in 8-day-old WT and *amiR-2572* seedlings, (n=18 for WT and 19 for *amiR-2572*). The color code corresponds to different LRP stages as indicated on the schematic root below. **f**Quantification of different LRP stages and emerged LR per root in (**a**), shown are averages (±SD), * indicates P < 0.05 (two-tailed Student’s t-test).

### Subgroup III ABCBs define a new group of plasma membrane-localized auxin exporters

Because multiple ABCBs have known auxin transport activities^26, 30, 31,^ we postulated that the targeted ABCB subfamily are also *bona fide* auxin transporters. Therefore, the coding regions of *ABCB15*, *B16*, *B17*, *B18* and *B22* were fused to *YFP*. Stable transgenic Arabidopsis YFP fusion lines were generated, all showing plasma membrane localization (Fig. 3a). Protoplasts prepared from Agrobacterium-transfected in *N. benthamiana* leaves showed increased IAA export for all 5 ABCBs at rates comparable to those of the canonical auxin transporting ABCB1^32^ (Fig. 3b). In contrast, none of these ABCBs enhanced export of the diffusion control, benzoic acid, (Fig. 3c). Together, the results suggest that the subgroup III ABCBs are plasma membrane localized auxin exporters. In agreement, all 5 ABCBs contained a conserved D/E-P motif in the C-terminal nucleotide binding fold that is diagnostic for auxin-transporting ABCBs^21^.

**Fig. 3:**
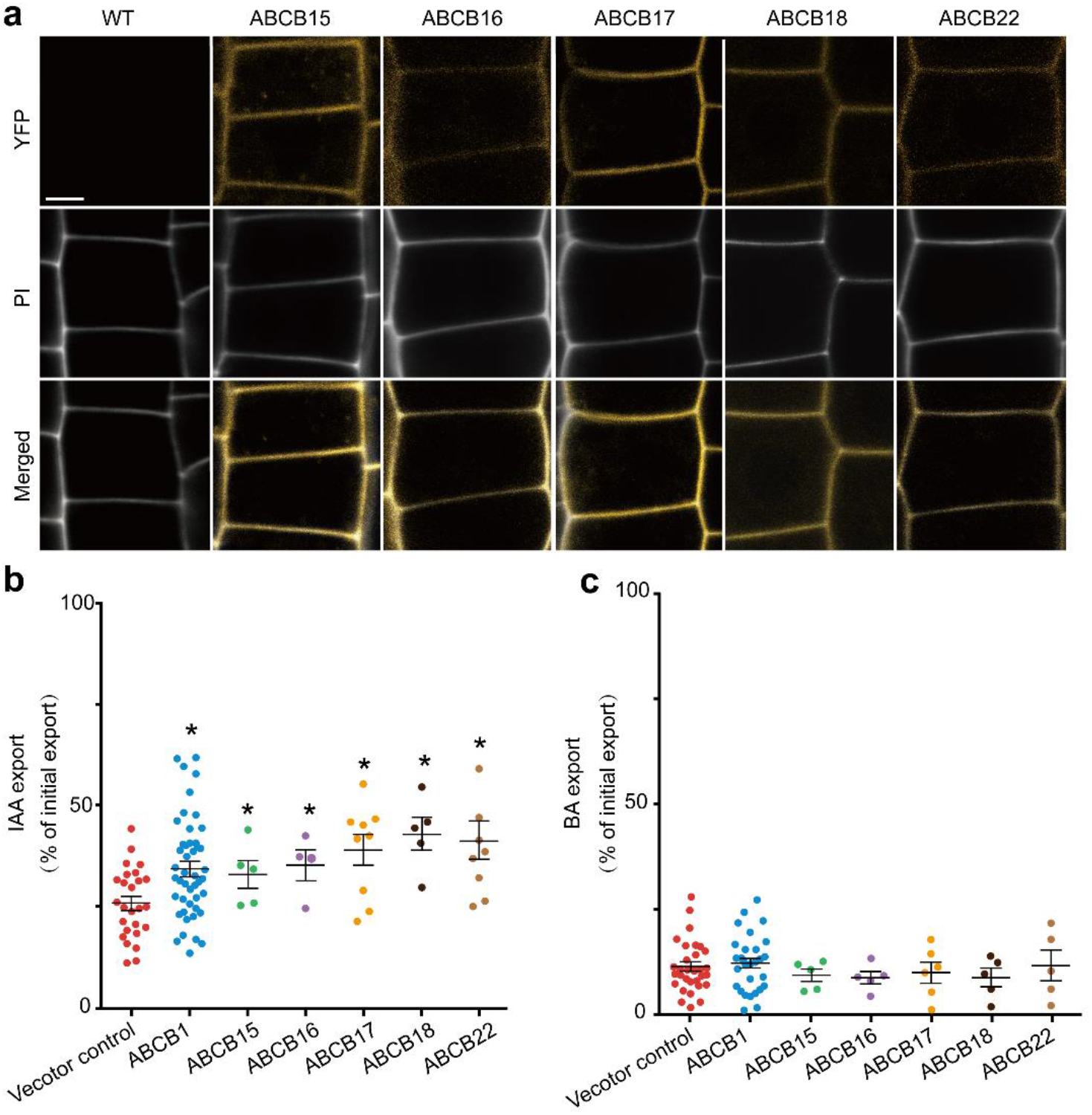
Subgroup III ABCBs localize to the plasma membrane and promote auxin transport. **a** Epidermal YFP fluorescence in primary root meristems of 3-day-old WT, *pro35S:YFP-ABCB15*, *pro35S:YFP-ABCB16*, *pro35S:YFP-ABCB17*, *pro35S:YFP-ABCB18*, *pro35S:YFP-ABCB22*. Cell walls are stained by Propidium Iodide (PI) in grey. Scale bars represent 10 μm. All pictures were analyzed using the same magnification. **b, c** IAA and BA export assay. Export of radiolabeled IAA (**b**) and benzoic acid BA (**c**) assayed in parallel from tobacco mesophyll protoplasts expressing indicated *ABCB*s of subgroup III against vector control. * indicates P < 0.05 (unpaired t-test with Welch’s correction) (mean ± SE; n ≥ 4 transport experiments generated from independent tobacco transfections).

Next, we analyzed the expression pattern of the five subgroup III ABCBs. We found strong expression of *ABCB15*, *B16*, *B17* and *B22* in the root meristem (Fig. 4a-d) and for *ABCB16* in all stages of LR development (Supplementary Fig. 6a). *ABCB18* was almost not expressed in the root meristem, but showed very weak expression in vascular tissues of the hypocotyl and mature root tissues (Fig. 4e; Supplementary Fig. 6c). Besides their root expression, *ABCB15*, *B17* and *B22* were also expressed in cotyledons, leaves and shoot meristem regions (Fig. 4a-d). Interestingly, detailed inspection of the expression pattern using confocal microscopy showed that their root meristematic expression was largely specific to the epidermis and LRC (Fig. 4e,f). This is consistent with a role in the auxin transport mechanism through which the LRC communicates with the main root for LR spacing.

**Fig. 4:**
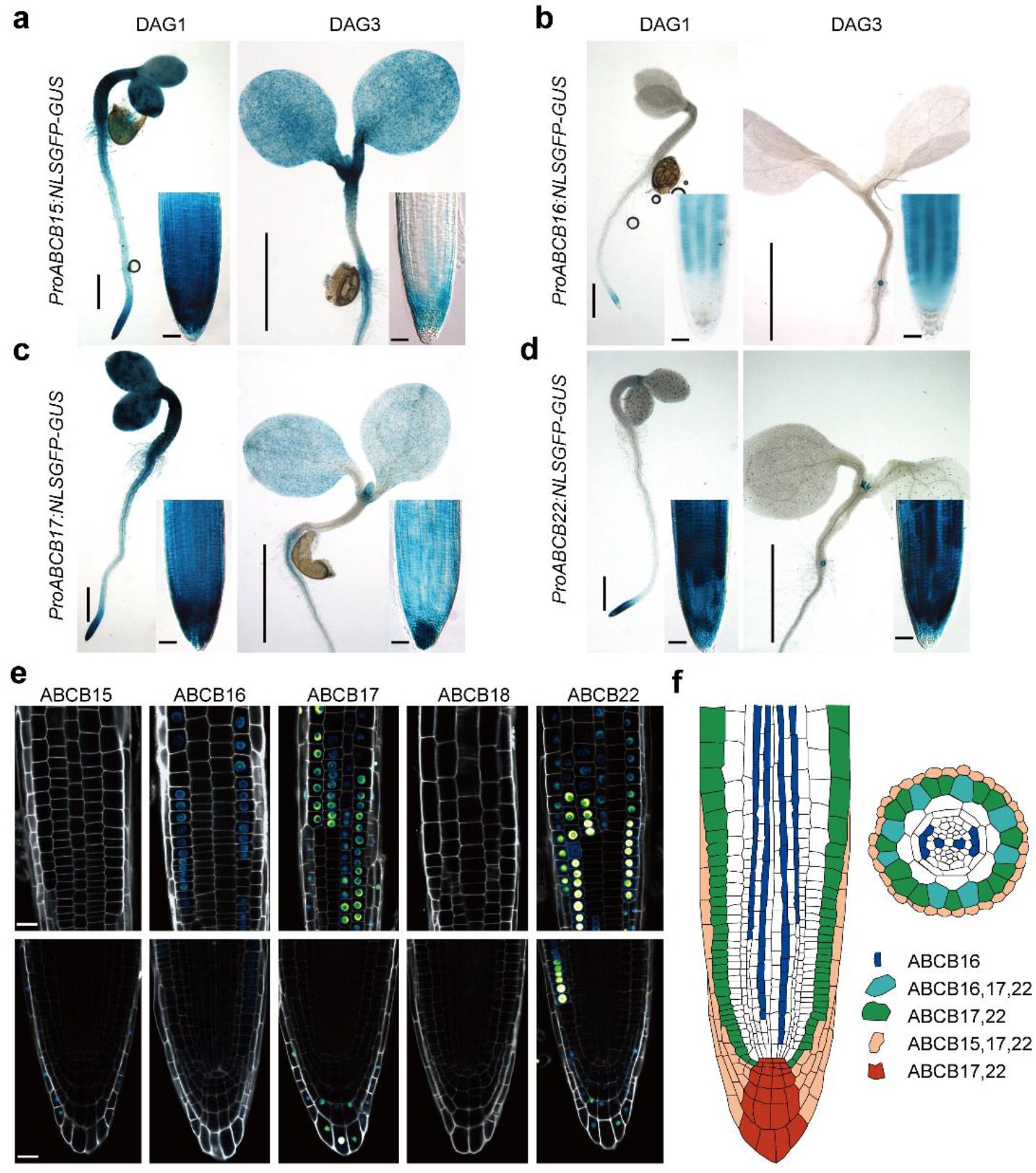
Overview of the expression patterns of subgroup III *ABCB*s. **a,b,c, d** *GUS* expression pattern of *proABCB_(15,16,17,18,22)_:NLSGFP-GUS* in 1- and 3-day-after germination seedlings (DAG). Scale bars = 0.5mm, for inset = 20 μm. **e**Surface and median view of GFP fluorescence in *proABCB15:NLSGFP-GUS*, *proABCB16:NLSGFP-GUS*, *proABCB17:NLSGFP-GUS*, *proABCB18:NLSGFP-GUS*, and *proABCB22:NLSGFP-GUS* expression in roots of 3-day-old seedlings. Propidium iodide in grey. Scale bars represent 20 μm. **f** Summary of *ABCB* subgroup III expression domains in the root apical meristem indicated on a longitudinal and radial section.

### Subgroup III ABCBs control the amplitude of auxin oscillations that instruct LR spacing

To test the hypothesis that subgroup III ABCBs control LR initiation, we zoomed into the origin of the LR defects of *AtTAS1c-ABCB-IIIa*. For two independent lines, we found that the reduced density of LRs was associated with an overall reduction in the number of LR primordia (LRP). Mainly, the early LRP stages (stages I, II and III) were reduced, without accumulating intermediate stages of LRP development (stages V-VII) (Fig. 5a,b). Similarly, *amiR-2572* showed a strong reduction in the early LRP stages, without accumulating intermediate LRP stages (Fig. 2e,f). These data demonstrate that the reduced density of emerged LRs in these lines is due to a defect at the level of initiation.

**Figure 5:**
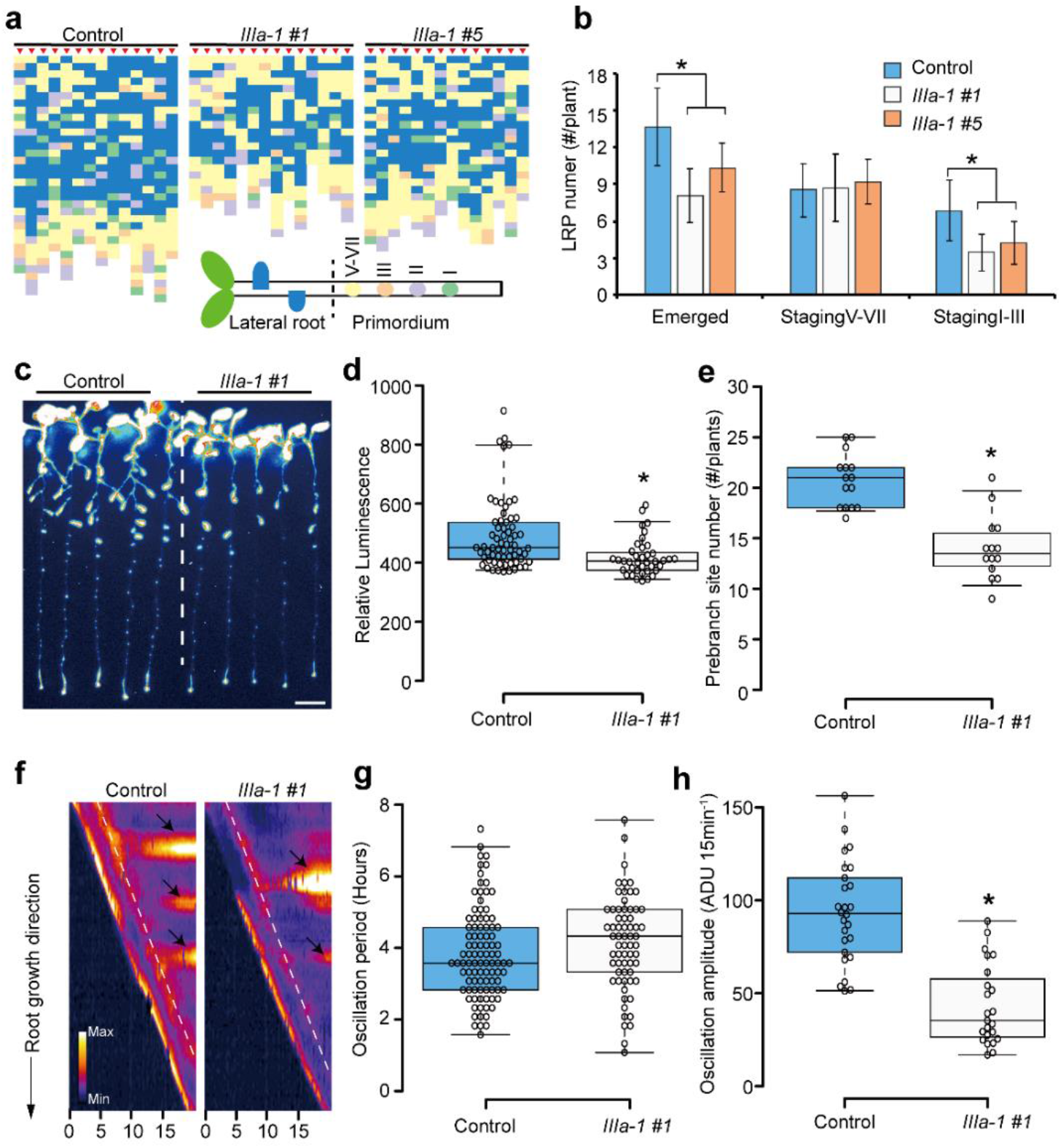
Subgroup III ABCBs control lateral root spacing by regulating pre-branch site number and the auxin oscillation amplitude. **a** Schematic representation of the distribution of emerged LRs and LRP along the primary root in 8-day-old WT and *AtTAS1c-IIIa* (Line1 and line5) seedlings (n = 10). The color code corresponds to different LRP stages as indicated in the schematic root. Each column represents an individual root indicated by red arrowheads. **b** Quantification of different LR stages and emerged LRs in (**a**), shown are averages (±SD) (n = 10). **c** Pre-branch sites in 8-day-old WT and *AtTAS1c-IIIa* line1 as determined by *DR5:LUC* luminescence. Scale bar represent 0.8 cm. **d** Quantification of *DR5:LUC* luminescence intensity of the pre-branch sites in 8-day-old WT (n = 64) and *AtTAS1c-IIIa* line1 (IIIa-1 #1; n = 39) seedlings. **e** Quantification of pre-branch site number per root in (**c**). n = 15 (WT), 14 (IIIa-1 line 1). **f** Kymograph of *DR5:LUC* intensity along the primary root in 3-day-old WT and *AtTAS1c-IIIa* line1 seedlings over 20 hr. *DR5:LUC* luminescence intensity is color coded (see color code in the bottom left corner of the panels) and plotted following the primary root elongation (y-axis) and time (x-axis). The arrows highlight pre-branch sites and white dashed lines indicate the *DR5:LUC* signal in OZ. **g, h** Quantification of the oscillation period (**g**) and amplitude (**h**) of *DR5:LUC* in 3-day-old WT and *AtTAS1c-IIIa* (line1) (n = 24). Center lines in box plots show the medians; box limits indicate the 25th and 75th percentiles as determined by R software; whiskers extend to 5th and 95th percentiles, outliers are represented by dots; data points are plotted as open circles. * indicates P < 0.05 (two-tailed Student’s t-test)

Lateral root initiation (stage I) is the first morphological hallmark of LR formation, and is preceded by a local maximum of auxin signaling, that can be visualized using *DR5:LUC*, referred to as prebranch sites^5^. Consistently with the reduced LR initiation, *AtTAS1c-IIIa* showed fewer prebranch sites (Fig. 5c-e). The *DR5:LUC* oscillation period in the *AtTAS1c-IIIa* was similar to that of the wild type (Fig. 5f,g), suggesting that LRC programmed cell death was not affected. In contrast, the *DR5:LUC* oscillation amplitude in the *AtTAS1c-IIIa* lines was significantly lower than in the WT (Fig. 5f,h), suggesting that not every auxin oscillation develops into a prebranch site and subsequently into a new LR. This suggests that subgroup III ABCB are part of the auxin homeostasis mechanism that determine *DR5:LUC* oscillation amplitude.

Previously, we inferred from tissue-specific complementation assays and *in silico* modeling that shootward auxin transport in the LRC is a critical determinant of oscillation amplitude^7, 11, 12^. The expression patterns of the subgroup III ABCBs and their knock-down phenotypes indicate that they could represent the elusive efflux component in this model. To test this hypothesis, we generated an estradiol-inducible auxin biosynthesis system driven specifically in the quiescence center (QC) (*pWOX5:XVE>>YUC1-2A-TAA1*). Simultaneous expression of *YUC1* and *TAA1* results in IAA synthesis from tryptophan^33, 34^. In the control, estradiol treatment induced a strong ectopic *DR5:VENUS* expression in the LRC, epidermis and the stele in the elongation zone within 7.5h. These effects were further enhanced after 9h estradiol treatment (Fig. 6a). This suggests that the auxin that was produced in the QC, was transported shootward via the LRC and epidermis towards the tissues of the elongation zone, where it activated *DR5-VENUS* expression. In *amiR-2572*, the induction of *DR5:VENUS* in the elongation zone was at both time-points severely reduced compared to the WT (Fig. 6a,b). These data demonstrate that subgroup III ABCBs contribute to shootward auxin transport in the meristem. These data, together with the expression patterns and phenotypes, are consistent with subgroup III-ABCBs being part of the shootward auxin flux in the outer layers of the meristem, that contributes to the *DR5:LUC* oscillation amplitude and LR spacing.

**Figure 6.**
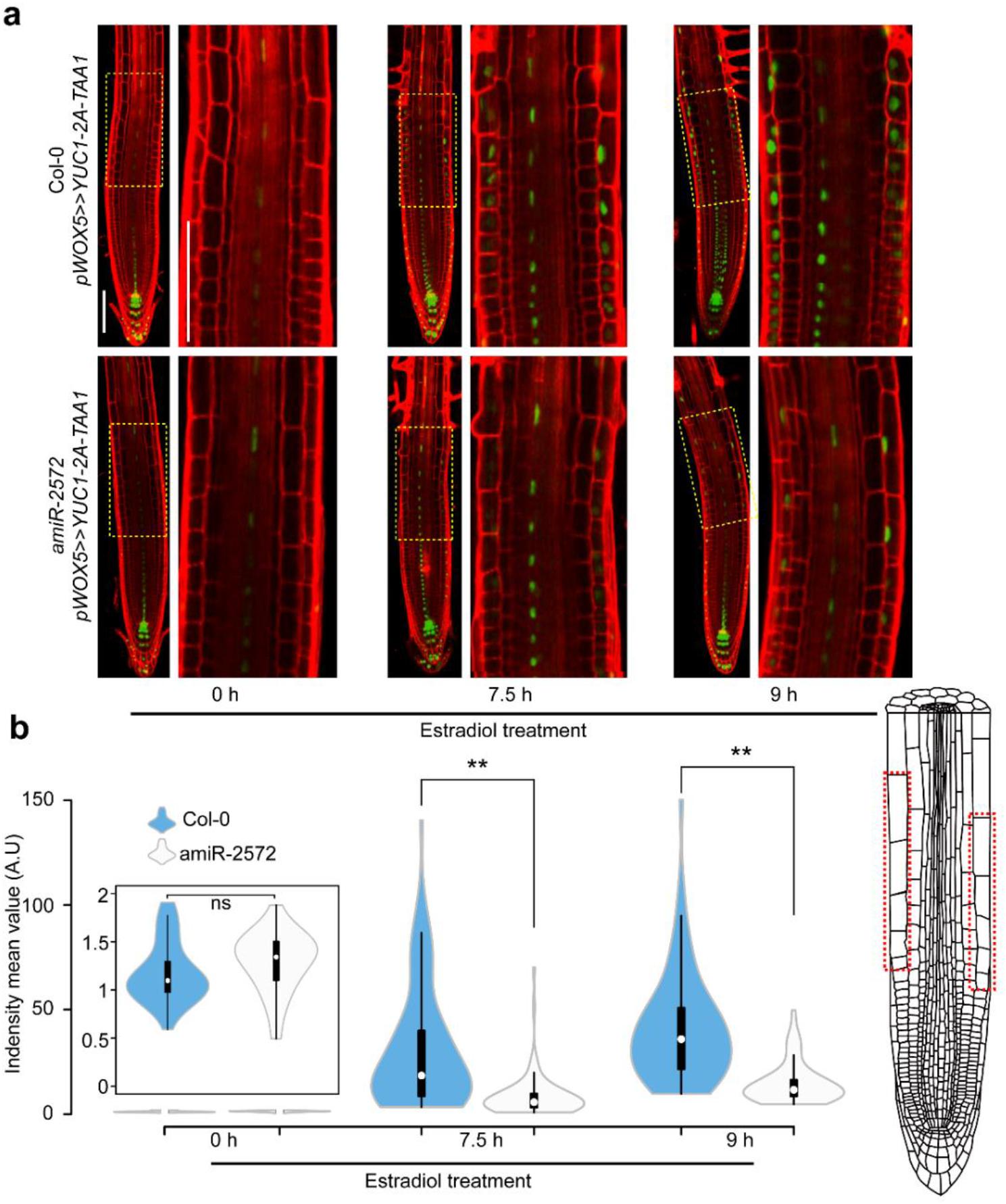
Subgroup III ABCBs contribute to shootward auxin transport in the LRC and meristem. **a** Analysis of *DR5:VENUS* expression in the root elongation zone of 4-day-old *pWOX5>>YUC1-2A-TAA1*, in Col-0 and *amiR-2572* treated with β-estradiol (5 μM) for 0, 7.5 and 9h. Propidium iodide in red. Yellow squares indicate zoomed pictures. Scale bar = 100 μM. **b** Quantification of *DR5:VENUS* signals in the epidermis of the elongation zone, as indicated in the scheme (Red square). White dots indicate the medians; box limits indicate the 25th and 75th percentiles as determined by R software; whiskers extend 1.5 times the interquartile range from the 25th and 75th percentiles; polygons represent density estimates of data and extend to extreme values.10 cells in the elongation zone per seedling and at least 4 seedlings of each treatment were measured ** indicates P < 0.01 (two-tailed Student’s t-test).

## Discussion

At its core, spacing of LRs in *Arabidopsis* can be simplified as the periodic activation of auxin signaling in the pericycle^5, 6, 12^. This model of LR spacing, assumes a local build-up of auxin that triggers LR initiation when an auxin signaling threshold is surpassed^12^. Surprisingly, the LRC plays a central role in this oscillatory auxin accumulation that determines LR spacing. On the one hand, the LRC contributes to the overall pool of auxin in the meristem via the local conversion of IBA to IAA^11^. On the other hand, dying LRC cells release auxin into the epidermis, resulting in a temporal rise in pericyclic auxin^7^. Periodic cell death in the LRC thus explains the oscillation of auxin activity in the pericycle to instruct LR spacing. In both cases, the LRC contribution to LR spacing assumes auxin transport from the LRC to the pericycle. This auxin transport mechanism involves the AUX1 IAA^−^:H^+^ uptake carrier in the LRC^7^. Currently, the molecular nature of the auxin efflux machinery involved in LR spacing remains elusive.

In an attempt to identify the missing auxin transporter(s), we delved deeper into the potentially massive functional redundancy within the ABCB gene-family, that contains multiple auxin transporters^30^. We identified a group of closely related *ABCBs* (*ABCB15*, *ABCB16*, *ABCB17*, *ABCB18* and *ABCB22*) that are required for LR spacing. We demonstrated that these ABCBs are plasma membrane-localized auxin exporters that contribute to the *DR5:LUC* oscillation amplitude, via effecting shootward auxin transport in the meristem. Interestingly, these *ABCB*s are largely co-expressed with *AUX1* in the outer tissues of the root meristem, suggesting they act in conjunction with AUX1 in the LRC and epidermis. Although this finding adds a new piece to the puzzle, it remains unclear whether this new cluster of ABCBs auxin transporter also bridges the cortex and endodermis, to feed into the pericyclic auxin pool and to surpass the critical auxin level that triggers LR initiation.

Recently, passive auxin diffusion via plasmodesmata, intercellular pores that linking the cytoplasm of adjacent cells, was shown to markedly improve the accuracy of simulated auxin distribution patterns in the root apical meristem^35^. Therefore, it will be of interest to evaluate the contribution of symplastic connectivity between the radial layers of the root to prebranch site formation and LR spacing.

## Materials and methods

### Plant material and growth conditions

*Arabidopsis thaliana* Colombia (Col-0) ecotype, was used as wild type. *abcb15-1* (SALK_034562), *abcb16-1* (SALK_006491), *abcb17-1* (SALK_002801), *abcb18-1* (SALK_013774), and *abcb22-1* (SALK_202270) mutant seeds were obtained from NASC. *Arabidopsis* transgenic lines *DR5rev:VENUS-N7*^36^ and *DR5:Luciferase* (*DR5:LUC*)^5^ were crossed and homozygous lines were selected and used as T0 for *AtTAS1c-ABCBs* transformation. Arabidopsis seeds were surface sterilized by chlorine gas, seeds were then sown in Petri dishes (12 cm X 12 cm) containing sterile half-strength Murashige and Skoog medium (0.5 × MS salts, 0.8% sucrose, 0.5 g/L 2-(N-morpholino) ethanesulfonic acid, pH 5.7, and 0.8% w/v agar), and grown under continuous light, after 3 days vernalization at 4℃.

### Plasmid construction

Most constructs were generated by the Gateway system^®^ (Invitrogen, Carlsbad, CA, USA). To construct the YFP fusion, coding sequences amplified from genomic DNA were cloned into pDONR-P2R-P3 (ThermoFisher Scientific) using the primers listed in Supplementary Table 1. The *35S* driven N-terminal YFP fusion expression clones were constructed by recombining pEN-L4-35S-R1^37,^ pEN-L1-Y-L2^37^ and the respective CDS clones into pH7m34GW using multisite LR Gateway reaction. For the promoter::NLSGFP-GUS reporters, ~2kb promoter fragments upstream of the coding sequence were amplified from genomic DNA using primers listed in Supplementary Table 2, and subsequently cloned into pENTR™ TOPO^®^ vector(pENTR™/D-TOPO^®^ Cloning Kits, ThermoFisher Scientific) to generate the corresponding entry clones. The promoter::NLSGFP-GUS was generated by performing an LR recombination reaction between Nuclear GFP fusion (pEN-L1-NF-L2) ^37,^ GUS reporter (pEN-R2-S*-L3)^37^ and pH7m34GW^37^. *AtTAS1c-ABCBs* constructs were generated using primers TAS-Ia/b-F/R, TAS-IIa/b-F/R and TAS-IIIa/b-F/R (Supplementary Table 3) as described^22^.

The *pWOX5:XVE>>YUC1-2A-TAA1* construct was generated by cloning the *YUC1-2A-TAA1* cassette into XhoI and SpeI sites of the pER8 vector^38^. The full-length cDNA of *YUC1* was cloned into the BamHI site and the full-length cDNA of TAA1 into the BglII site of the pM2A vector containing 2A peptides^39^. For QC-specific activation of the *YUC1-2A-TAA1* cassette, the genomic DNA of *WOX5* promoter (WOX5pF: CAATATATCCTGTCAAACaaagacttttatctaccaacttcaa; WOX5pR: GCCGTTAACGCTTTCATcgttcagatgtaaagtcctcaactgt) was used.

### Generation of *pWOX5:XVE>>YUC1-TAA* lines

*pWOX5:XVE>>YUC1-2A-TAA1* (*pWOX5>>YUC1-TAA1*) was introduced into DR5:VENUS background by transformation and 10 independent lines were selected. Homozygous lines for both *pWOX5>>YUC1-TAA1* and *DR5:VENUS* were crossed to amiR-2572 lines to generate F1 seeds. Homozygous plants for *pWOX5>>YUC1-TAA1*, *DR5:VENUS* and *amiR-2572* were gained by resistance selection and phenotyping in F3 population. Similarly, amiR-2572 line was crossed with *DR5:VENUS* to generate F1 seeds and F3 homozygous for both constructs were obtained.

### *Agrobacterium* and *Arabidopsis* transformation

*Agrobacterium tumefaciens* strain GV3101 was transformed with the relevant binary plasmids via the freeze-thaw procedure^40^. An individual PCR confirmed Agrobacterium colony was used for floral dip^41^. Transformants were selected and the segregation of the T2 analyzed using appropriate antibiotics.

### Phenotyping and LR staging

To quantify the LR phenotype in wild-type plants and mutants, emerged LR of whole seedlings were counted under a dissecting microscope, 8 days after germination. Root lengths were measured via Fiji (ImageJ 1.52n^42^) using digital images obtained by scanning the Petri dishes.

To analyze the LR primordium stages, root samples were cleared as described previously^43^. All samples were analyzed by differential interference contrast microscopy (Olympus BX51).

### Oscillation and prebranch site

The Luciferase imaging of whole seedlings and oscillation expression analysis was performed as described^44^. A Lumazon FA imaging system (Nippon Roper) carrying a CCD camera from Princeton Instruments Ltd. (Trenton, NJ, USA) or NightSHADE LB985 in vivo plant imaging system (BERTHOLD TECHNOLOGIES) carrying a deep-cooled slow scan CCD camera from Andor Instruments Ltd. (Belfast, UK) were used for luciferase imaging.

To monitor the pre-branch site numbers, we used 8-day-old DR5:LUC seedlings for pre-branch site quantification. The D-luciferin solution (1 mM) was sprayed gently on the seedlings, and kept for 10min in the dark and imaged in the Lumazon system with a 15-minute exposure time.

For Long-Term Imaging of Luciferase Signal in the root tip, square plates containing 1/2MS medium were sprayed with 1mM D-Luciferin solution (0.01% Tween80) and left to dry in the dark. Then 3-day-old DR5:LUC seedlings were transferred on the plates and imaged immediately with a macro lens every 10 minutes with a 7-minute exposure time for indicated times. The period of the *DR5:LUC* oscillations was determined based on the number of frames that spaced a DR5:LUC maximum in the OZ of each seedling root, multiplied with the time of each cycle.

### Kymograph

Kymographs (http://www.embl.de/eamnet/html/body_kymograph.html) were generated by ImageJ to visualize the spatiotemporal changes of DR5:LUC signal in the root tips during primary root growth. For this purpose, a time-lapse movie (TIFF series) was loaded into ImageJ, and a “Z-projection” was performed to have an overview of the luciferase signal changes following primary root growth over time. Subsequently, a segmented line was drawn on the newly formed primary root and marked by the “ROI manage” function. This line was restored in the original TIFF series to generate “MultipleKymograph”. In our experiments, 3-day-old seedlings were used.

### Confocal microscopy

For reporter lines and translational fusion, seedlings were imaged on a Zeiss 710 confocal microscope. For the propidium iodide (PI)-treated root images, seedlings were stained with 2 μg/mL PI for 3 minutes, washed with water, and used for confocal imaging. For root imaging, GFP was excited at 488 nm and acquired at 500 to 530 nm. YFP was excited at 514 and the emission between 519-564 nm was collected for YFP and between 614-735 nm for PI.

For the *pWOX5>>YUC1-TAA* experiments seeds were sown on MS plates, stratified at 4°C for 2 days, and grown vertically in growth chamber for 4 days at 21°C. 4-day-old seedlings of the *pWOX5:YUC1-TAA1*, *DR5:VENUS* in Col-0 and *amiR-2572* background were treated with 5 μM estradiol for the indicated time-points. Seedlings were stained in 10 mg L-1 propidium iodide for 2 min and rinsed in water for 30 s. Confocal microscopy was performed using a Zeiss LSM780 inverted confocal microscope equipped with a 20×/0.8 M27 objective lens. VENUS and propidium iodide were excited using an argon-ion laser and a diode laser, respectively. VENUS was excited at 514 nm and detected at 518-588 nm, propidium iodide was excited at 561 nm and detected at 588-718 nm.

### GUS staining and root sectioning

The GUS assay was performed as previously described^45^. For Arabidopsis cross-section root specimens, GUS stained seedlings were subjected to fixation, dehydration and embedding as previously described^46^. GUS-stained tissues were imaged using a Leica Bino and Olympus BX51 microscope for different tissues.

### Construction of phylogenetic tree

The evolutionary history was inferred using the Neighbor-Joining method^47^. The percentage of replicate trees in which the associated taxa clustered together in the bootstrap test with 1000 replicates^48^. The tree is drawn to scale, with branch lengths in the same units as those of the evolutionary distances used to infer the phylogenetic tree. The evolutionary distances were computed using the Maximum Composite Likelihood method and are in the units of the number of base substitutions per site. The analysis involved 21 nucleotide sequences. All positions containing gaps and missing data were eliminated. There were a total of 3455 positions in the final dataset. Evolutionary analyses were conducted in MEGA7^49^ and optimized via Interaction Tree Of Life (https://itol.embl.de/).

### Genotyping

T-DNA lines for the ABCB single mutants were ordered from The *Arabidopsis* Information Resource (https://www.arabidopsis.org/), and genotyping primers for T-DNA insertion were designed using the T-DNA Primer Design Tool powered by Genome Express Browser Server (GEBD) (http://signal.salk.edu/tdnaprimers.2.html). Homozygous mutants were selected by PCR performed with primers listed in Supplementary Table 4.

### RNA extraction and Nanostring

Total RNA was isolated from the indicated plant materials using the PureLink RNA Mini Kit (Invitrogen). Nanostring transcript quantification was done as described previously^26^.

### Auxin transport measurements

Simultaneous ^3^H-IAA and ^14^C-benzoic acid (BA) export from tobacco (*N. benthamiana*) mesophyll protoplasts was analyzed as described^19^. Tobacco mesophyll protoplasts were prepared 4 days after agrobacterium-mediated transfection with *proS35S:ABCB1-YFP, pro35S:YFP-ABCB15, pro35S:YFP-ABCB16, pro35S:YFP-ABCB17, pro35S:YFP-ABCB18, pro35S:YFP-ABCB22.* Relative export from protoplasts is calculated from exported radioactivity into the supernatant as follows: (radioactivity in the protoplasts at time t = 10 min.) - (radioactivity in the supernatant at time t = 0)) * (100%)/ (radioactivity in the supernatant at t = 0 min.); presented are mean values from >4 independent transfections.

### CRISPR/Cas9 mutagenesis and selection of mutant alleles

Four single-guide (sg) RNAs were designed using the CRISPR-P tool (http://cbi.hzau.edu.cn/cgi-bin/CRISPR)^50^ to align the ABCBs coding sequence. The sgRNAs are designed to target multiple ABCBs at once: sgRNA-19 targets ABCB16, 18, 22, 17 (20% cleavage), and 15 (0.3% cleavage); sgRNA-20 targets ABCB18, 22, 16, (92% cleavage), 17 (10% cleavage) and 15 (0.1% cleavage); sgRNA3 targets ABCB16, 18, 17 (0.4% cleavage), and 15 (0.1% cleavage) and sgRNA4 targets ABCB16, 17, 18 (49% cleavage), and 15 (0.1% cleavage). Vectors were assembled using the Golden Gate cloning system^51^. The sgRNA-19, sgRNA-20, sgRNA-3 and sgRNA-4 were cloned downstream of the *Arabidopsis* U6 promoter (pATU6) in the Level 1 acceptors pICH47761, pICH47772, pICH47781 and pICH47791, respectively, as previously described^52^. The Level 1 constructs were assembled in the binary Level 2 vector pAGM4723. sgRNA sequences are listed in Supplementary Table 5. Genotyping was carried out using primers listed in Supplementary Table 6.

For the large fragment deletion mutant (*ldf*), 3 sgRNAs were designed targeting ABCB15, and another 3 sgRNAs were designed targeting ABCB22. The sgRNAs are listed in Supplementary Table 7. Six sgRNAs were assembled into *pFASTRK* as described^53^. Pooled T1 plants were screened for a 0.5-1kb amplicon, using primers spanning the 54kb genomic fragment, as an indicator of the deletion. In the T2 generation, individuals lacking the Cas9 transgene were screened for the amplicon, which was sent for Sanger sequencing. Genotyping was done using primers listed in Supplementary Table 8.

In order to confirm the correct excision and deletion of the targeted region from the genome, we used NGS. Sequencing was perfomed on an illumina HiSeq 4000 machine, which yielded 47,827,766 reads (150nt PE), being 57.8x coverage. The reads were then aligned to the reference ATH Col-0 genome using BBMap (https://jgi.doe.gov/data-and-tools/bbtools/) using default settings. The obtained BAM files containing the aligned reads was subsequently processed with bedtools genomecov^54^ (parameter settings: -bga −split). This resulted in a coverage plot reflecting the sequencing depth over the ATH genome sequence. Exploring the coverage plot clearly showed that the targeted regions was no longer present (indicated by having a coverage of zero), in our re-sequenced line. Moreover, it also showed no off-target modifications, nor that the excised region would have been reinserted elsewhere in the genome.

Data supporting the NGS analysis part of this study has been deposited at the ENA under BioProject number: PRJEB38980.

## Acknowledgements

We thank Jose Alonso and Thomas Jacobs for providing early access to unpublished materials at the beginning of this project. This work was supported by grants from the Swiss National Funds (31003A-165877/1) to M.G., the China Scholarship Council to J.C., the European Research Council Starting Grant (757683-RobustHormoneTrans) to E.S., the PBC postdoc fellowship to Y.H and Y.Z.

